# Mapping the architecture of regulatory variation provides insights into the evolution of complex traits

**DOI:** 10.1101/2020.05.24.113217

**Authors:** Offir Lupo, Gat Krieger, Felix Jonas, Naama Barkai

## Abstract

**Background:** Organisms evolve complex traits by recruiting existing programs to new contexts, referred as co-option. Within a species, single upstream regulators can trigger full differentiation programs. Distinguishing whether co-option of differentiation programs results from variation in single regulator, or in multiple genes, is key for understanding how complex traits evolve. As an experimentally accessible model for studying this question we turned to budding yeast, where a differentiation program (filamentous) is activated in *S. cerevisiae* only upon starvation, but used by the related species *S. paradoxus* also in rich conditions.

**Results:** To define expression variations associated with species-specific activation of the filamentous program, we profiled the transcriptome of *S. cerevisiae*, S. *paradoxus* and their hybrid along two cell cycles at 5-minutes resolution. As expected in cases of co-option, expression of oscillating genes varies between the species in synchrony with their growth phenotypes and was dominated by upstream *trans-*variations. Focusing on regulators of filamentous growth, we identified gene-linked variations (*cis*) in multiple genes across regulatory layers, which propagated to affect expression of target genes, as well as binding specificities of downstream transcription factor. Unexpectedly, variations in regulators essential for *S. cerevisiae* filamentation were individually too weak to explain activation of this program in *S. paradoxus*.

**Conclusions:** Our study reveals the complex architecture of regulatory variation associated with species-specific use of a differentiation program. Based on these results, we suggest a new model in which evolutionary co-option of complex traits is stabilized in a distributed manner through multiple weak-effect variations accumulating throughout the regulatory network.

## Introduction

New phenotypes arise in evolution from mutations that change gene function or regulation. At the molecular level, the majority of mutations are neutral or of small phenotypic effects[1]. Still, evolutionary related species that express the same set of proteins differ in complex phenotypes, including size, growth pattern, and body morphology[2–4]. A compelling model is that complex traits do not evolve *de-novo* but emerge through recruitment (co-option) of existing gene expression program to new contexts. Evidence supporting co-option were presented in the context of morphological evolution, where major interspecies differences in the positioning of body appendages or wing patterns were linked to variations in *cis*-regulatory elements controlling the expression of major regulators, including homeobox transcription factors or developmental signaling proteins[5–10].

Modulating the expression of single upstream regulators provides a genetic shortcut for the evolution of complex traits[11]. Complicating this view, however, is the fact that gene expression networks, such as these activating complex traits, are polygenic and include multiple activators and inhibitors that could act as drivers[12]. Further, due to the inter-connected nature of genetic circuits, variations in seemingly distant genes could propagate to influence the same phenotype[13]. This was exemplified recently in *Drosophila*, where expression variation in a Hox gene that correlated with morphological differences was found to be of little phenotypic consequence due to its masking by variations in other genes within the same pathway[14]. Understanding the genetic basis of complex traits therefore requires expanding our view from single regulators to full genetic circuits. In particular, it raises the question of whether co-option of regulatory programs result from variation in a single upstream regulator, or from multiple variations distributed among different genes. This question is difficult to address using the existing multicellular models of co-option, whose complexity precludes systematic analysis of all regulators.

As a model to address this question, we turned to budding yeast, which provides an experimentally accessible model for revealing the architecture of genetic variations associated with phenotypic differences. In multicellular models, co-option refers to the recruitment of existing differentiation programs to a new spatial-temporal context [15,16]. A unicellular analog of this process is the recruitment of existing differentiation programs to a new set of environmental conditions. Budding yeast undergo a dimorphic transition between yeast-form and filamentous growth modes, presenting two alternative growth programs which differ in multiple properties, including the selected budding site, cell morphology and adhesion, and the relative lengths of the different cell cycle phases[17]. In the model yeast *Saccharomyces cerevisiae*, transition to filamentous growth occurs mostly in nutrient-poor environments, allowing non-motile cells to forage for nutrients[17–19]. By contrast, *S. paradoxus* (CBS432), a close relative of *S. cerevisiae*, becomes filamentous even in rich media[20,21]. Therefore, the two species activate distinct differentiation programs when presented with the same condition, providing a model for evolutionary recruitment.

In addition to allowing systematic and rapid analysis, the budding yeast model offers additional key advantages for studying the basis of evolutionary recruitment. First, studies in *S. cerevisiae* characterized the regulatory circuits triggering the yeast to filamentous growth transition, identifying hundreds of genes that affect it and can therefore act as potential drivers of this adapted-response[22–28]. Moreover, budding yeast can mate to form inter-specific hybrids. Profiling allele-specific expression in hybrids allows to systematically distinguish variations that are linked to the gene itself (*cis*) from the that propagate from upstream variations (*trans*), a key for defining the genetic basis of expression divergence[29–38]. This is because within the hybrid, the two genomes are subject to the same *trans* environment, so that differences in the expression of the two alleles must result from *cis* effects. Making this distinction is particularly relevant in the case of co-option, where the *cis-*variations driving recruitment are expected to lead to multitude of propagating *trans* effects.

Our goal in this study was to reveal the architecture of expression variation underlying the differences in growth phenotypes of *S. cerevisiae* and *S. paradoxus* when growing in the same rich conditions. To this end, we examined how genes and cellular processes involved in the yeast-to-filamentous transition are expressed in the two species. By profiling the cell-cycle transcription programs of the two species and the allele-specific expression within their hybrid, we found that central pathways regulating the yeast-to-filamentous transition accumulated *cis* effects that bias the expression of regulators in the direction favoring the selected program. These were found in the main signal transduction components, transcription factors (TFs) and their downstream target genes. Similarly, we found variations in cell cycle TFs that propagated through a combination of *cis* and *trans* effects to influence both the expression of target genes and promoter binding specificities of another TF. Single gene candidates, tested by either deletion or promoter-swap, were insufficient to fully account for the phenotypic inter-species differences on their own. Our results suggest a model whereby co-option of differentiation programs is stabilized through the distributed action of multiple small-effects acting at different steps of the multi-layered response.

## Results

### Evolutionary related budding yeast species employ distinct growth program when grown in the same condition

In rich conditions, *S. cerevisiae* cells grow in yeast-form. *S. paradoxus*, a species closely related to *S. cerevisiae*, appears filamentous when grown in those same conditions. Using live microscopy, we verified that *S. paradoxus* cells display the hallmarks of filamentous growth: increase in cell length, reorganization of budding polarity, lack of mother-daughter separation and enhanced cell to cell adhesion (Figs 1A and 1B).

**Fig 1.**
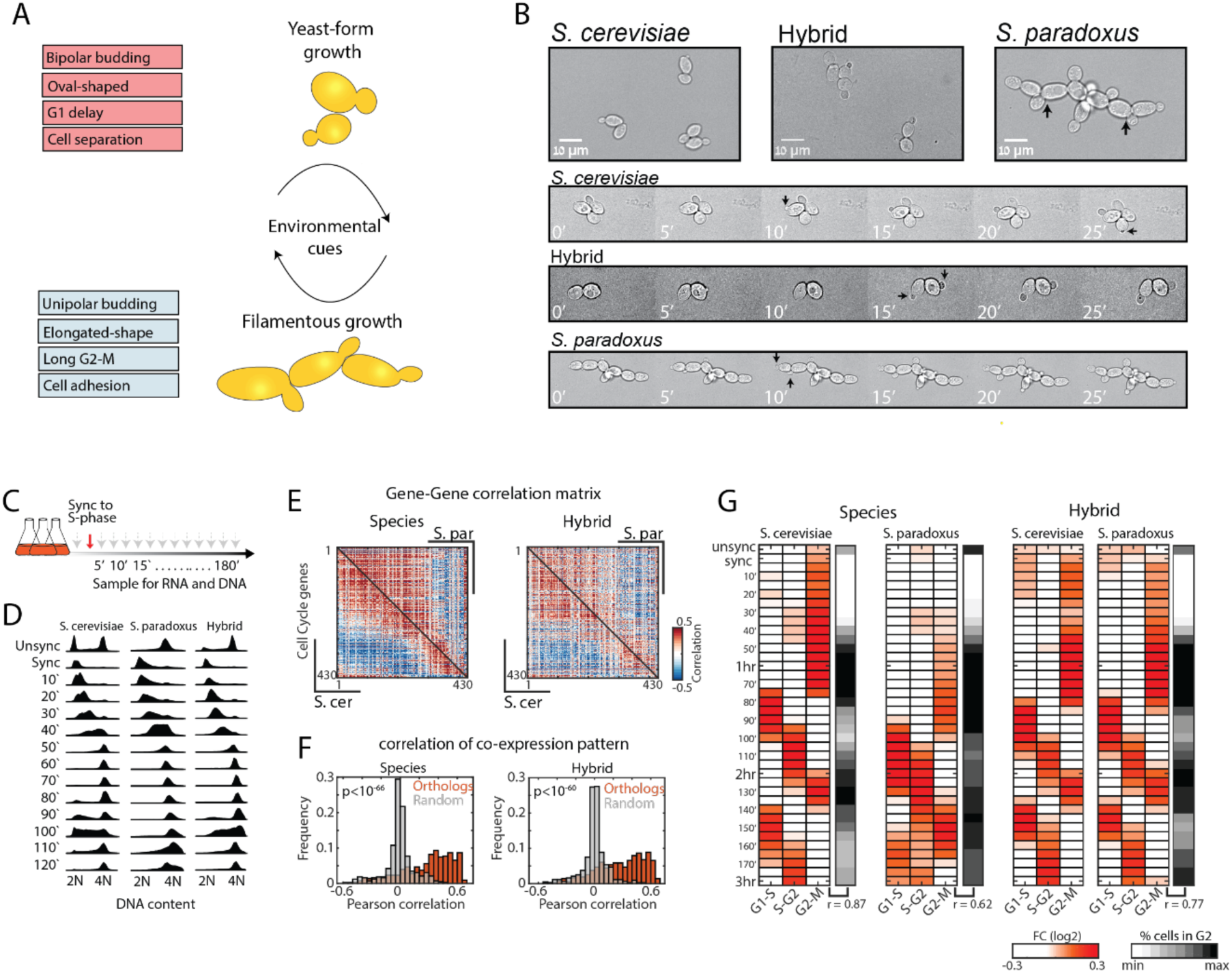
*Trans*-dominant expression program follows the growth pattern. **(A)** A scheme of the two types of growth in budding yeast. Budding yeast can undergo a dimorphic shift upon environmental cues which involves modifying different processes in cell cycle and morphology. **(B)** Microscopy images of *S. cerevisiae, S. paradoxus* and their hybrid grown on YPD. *S. cerevisiae* grows as yeast while *S. paradoxus* grows in filaments. The hybrid grows as yeast, while exhibiting short daughter-delay. Note cell elongation, unipolar budding and synchronized mother-daughter budding in *S. paradoxus*. Black arrows point at new buds. **(C)** Experimental layout: Cells were synchronized with HU, washed and sampled every 5 minutes for three hours. **(D)** Synchronized progression was monitored by measuring the DNA content using flow-cytometry (full profiles are shown in Fig S1A) **(E)** Co-expression of cycling genes is conserved between species. Shown are gene-gene correlation matrices (Pearson) of periodic genes ordered by their expression peak along the cell cycle, *S. cerevisiae* is shown in the lower triangle and *S. paradoxus* in the top triangle. **(F)** Histograms showing the distributions of co-expression correlation coefficients between orthologous genes and between random pairs (indicated p-value of a two-samples t-test). **(G)** Periodicity of gene expression is synchronized with cell-cycle transitions. Shown are the average fold changes of periodic genes (relative to median) classified to three groups based on expression time. Black panel indicated percent of cells with 4N DNA content. Correlation values in bottom indicate the Pearson correlation coefficient between expression of the G2-M module to percent cells with 4N DNA content.

The changes associated with transition between yeast-form and filamentous growth, such as cell shape, budding pattern and phases duration are inherently coupled to cell cycle regulation[17,18]. We therefore asked whether the gene expression program of the two species is coordinated with their distinct cell cycle program. We grew diploids of both species in rich media, arrested them in early S-phase using hydroxyurea (HU), and profiled their gene expression at five minutes intervals following release from the arrest for three hours, during which they underwent two full cell-cycles (Fig 1C). Synchronized progression of the cells was validated by measuring the DNA content (Figs 1D and S1A)

Previous studies in *S. cerevisiae* defined the set of genes showing periodic expression along the cell cycle[39–41]. We assembled this list of cell-cycle regulated genes, and examined their dynamics along our time courses. The majority of these genes were expressed periodically in both species (72% and 65% in *S. cerevisiae* and *S. paradoxus*, respectively, Figs S1B-D). Further, co-expression patterns between orthologs showed high correlations (Fig 1D), confirming an overall conserved cell cycle transcription program. Classifying the genes to groups based by their cell-cycle expression time, revealed periodicity that was in synchrony with cell-cycle phases, as defined by DNA content (Fig 1E). In particular, the G2-M delay and short G1 characteristics of the *S. paradoxus* cycle were evident in the expression of genes activated at these phases. For example, S-phase genes followed G1 genes at 10 minutes delay in *S. paradoxus*, compared to 20 minutes delay in *S. cerevisiae* (Fig 1E). Accordingly, similarity between expression profiles of the two species followed the cell cycle phase, rather than sampling time (S1E Fig).

The global synchronization of gene expression with the cell division cycle is indicative of a global dynamic guided by variations in upstream *trans* regulators. We tested this by profiling the interspecific hybrid, where both genomes are subject to the same *trans* environment. The hybrid grew as yeast, indicating the dominance of this program (Fig 1B). Consistent with previous report[20], it progressed faster through the cell cycle, showing 5% and 15% faster growth than *S. cerevisiae* and *S. paradoxus*, respectively. Expression of periodic genes within the hybrid was synchronized with cell-cycle progression, with the two genomes following the same dynamics (Fig 1E), therefore regulated in *trans.* Furthermore, comparing the expression levels of periodic genes revealed that *trans* effects dominate also variations in expression levels in addition to expression dynamics (S1F Fig). Therefore, the majority of variations in the expression of cell cycle genes propagates from variations in upstream acting *trans* factor(s).

### Cis-effects bias expression of yeast-to-filamentous regulators

In *S. cerevisiae*, ectopic expression or deletion of specific regulators can induce filamentous growth independently of growth conditions[17,18]. This includes changes in the main signaling pathways, cell cycle regulation or in other genes that have an effect on the filamentous growth phenotype (Fig 2A). We reasoned that variations in regulators expression could also account for the inter-specific differences in growth modes. To systematically search for such variations, we assembled lists of genes which deletions or over-expressions affect the yeast-filamentous transition in *S. cerevisiae*[42] (see methods). Comparing the average expression of these genes along our time course, we found a bias favoring the respective growth mode (S2A Fig). Specifically, genes that have a negative effect on filamentous growth were more abundant in *S. cerevisiae*, while those having a positive effect were more abundant in *S. paradoxus*. Surprisingly, this bias was maintained in the hybrid, indicating its *cis* origin.

**Fig 2.**
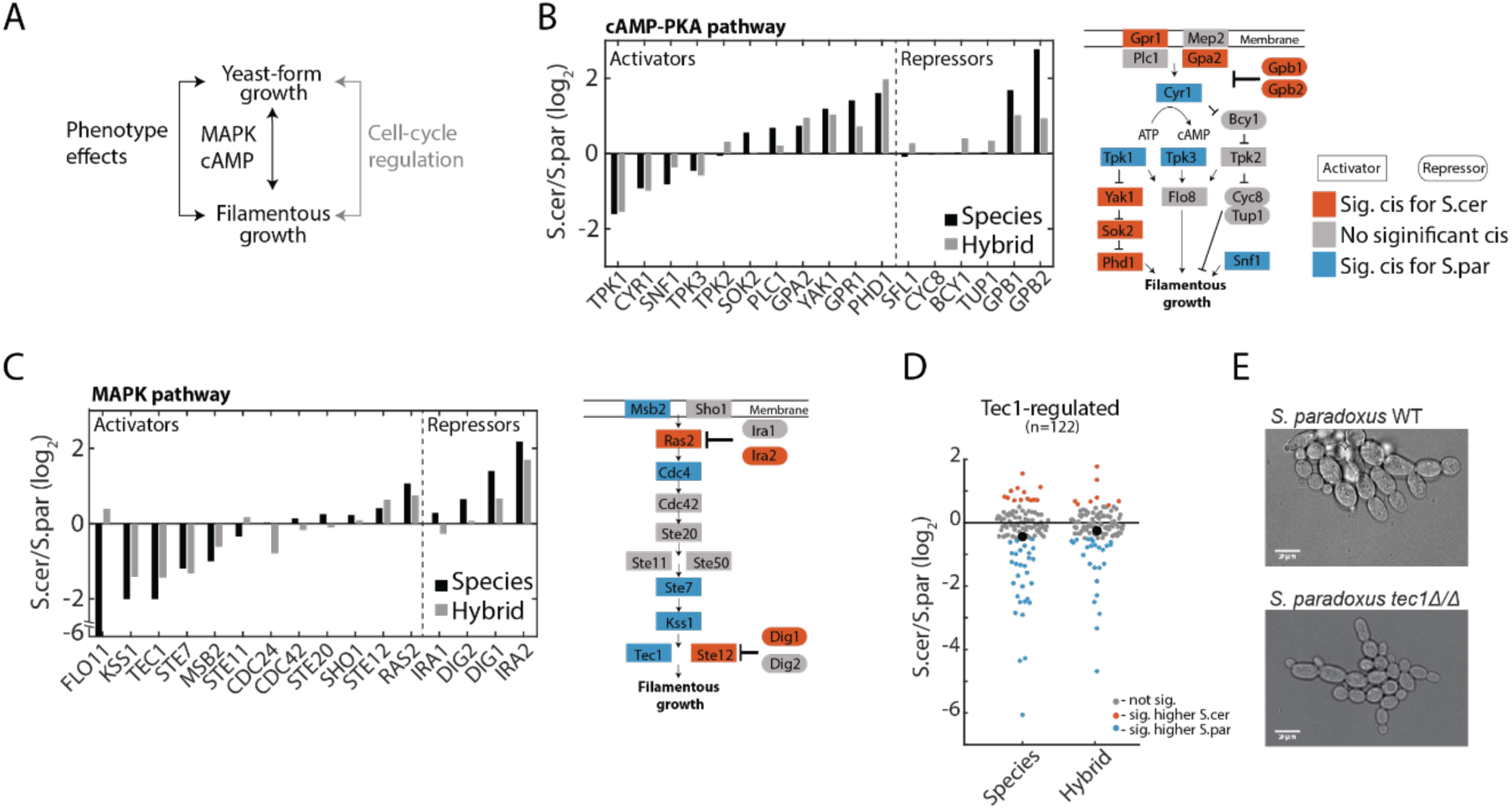
Filamentous regulators accumulated *cis-*effects that bias expression towards the selected growth program. **(A)** a scheme representing different regulatory layers that affect the yeast-to-filamentous growth switch. (**B-C)** Genes in the main filamentous signaling pathways cAMP-PKA (B) and MAPK (C) show differential expression in *cis* and in *trans*. Left: Shown are log_2_ fold change (FC) values of genes involved the indicated pathways. Right: Schemes representing the respected signaling cascades. Colors are given for genes showing significant FC difference between hybrid alleles (*cis*), as indicated in legend. **(D)** *cis*-regulation of Tec1-dependent genes. Shown are FC distributions of Tec1-regulated genes (Fig S2B). Red and blue dots indicate significantly higher expression in *S. cerevisiae* and *S. paradoxus*, respectively. Black dot marks the mean **(E)** Phenotypes of diploid *S. paradoxus* WT and *tec1*Δ/Δ, growing in rich conditions (YPD).

We next examined the two main signaling pathways that control filamentous growth, MAPK and cAMP-PKA. The MAPK cascade activates Tec1, the transcription factor (TF) acting as a master regulator of the filamentous program. The second pathway, the cAMP-PKA cascade, activates the TF Flo8 and inhibits the general co-repressors Tup1-Cyc8. In both pathways, repressors were expressed to higher levels in *S. cerevisiae* (Figs 5B and 5C; 5 out of the 5 differentially expressed repressors), while activators in MAPK pathways were more highly expressed in *S. paradoxus* (Fig 5C 5/6). Activators in the cAMP pathway did not show any bias (Fig 5B 4/10). While most expression variations were small, expression of TEC1 stood out as being 4-folds higher in *S. paradoxus*, while its inhibitors, DIG1 and DIG2, were expressed at higher levels in *S. cerevisiae* (Fig 5C; 2.6 and 1.5 folds, respectively). Further, comparing alleles expression within the hybrid revealed that in the majority of genes variations result from a combination of *cis* and *trans* effects, mostly acting in the same direction (18/20), promoting the selective phenotype.

**Fig 3.**
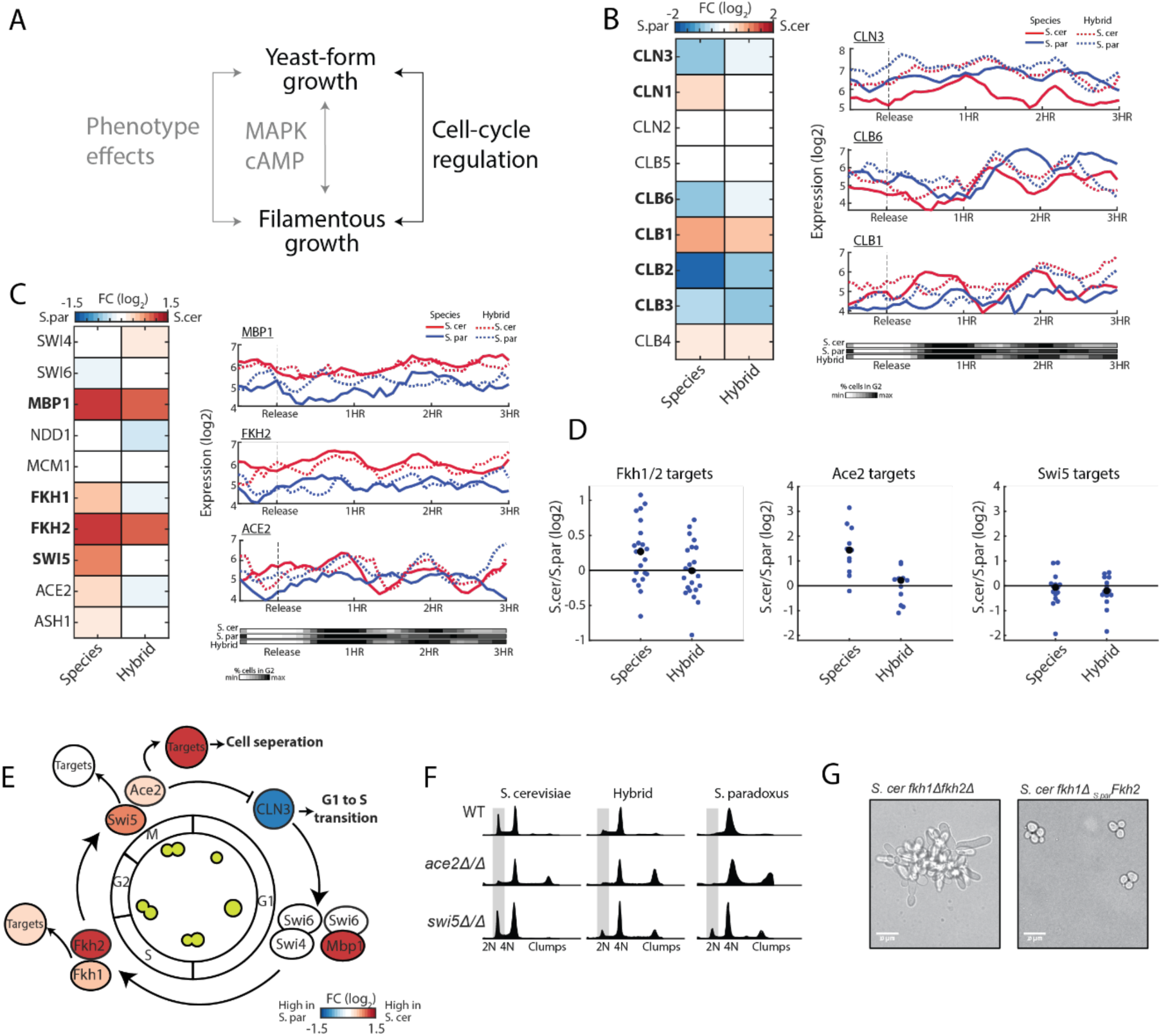
Cell-cycle regulators accumulated *cis-*effects that bias expression towards the selected growth program. **(A)** A scheme representing different regulatory layers that affect the yeast-to-filamentous growth switch. **(B)** Expression of cyclins. Left: Examples of CLN3 (early G1), CLB6 (early S) and CLB1(G2) along the cell cycle, in synchrony with changes in DNA content (bottom panel). Right: log_2_ FC expression differences. Genes ordered by their expression time along the cell cycle. Genes in bold have significant differential expression between species. **(C)** Expression of cell cycle transcription factors (TF), same as in (A). **(D)** log_2_ FC in Fkh1/2, Ace2 and Swi5’s targets. Number indicates number of genes; Black dots indicate the group’s mean. **(E)** Divergence in cell cycle regulators. A scheme of cell cycle regulation, color indicates inter-species expression differences. **(F)** DNA contents profiles of WT, *ace2*Δ/Δ and *swi5*Δ/Δ, in both species and hybrid, taken during exponential growth in YPD. Grey area marks cells with 2N DNA content (in G1 phase). **(G)** Phenotype of *S. cerevisiae fkh1*Δ *fkh2*Δ (left) and *S. cerevisiae fkh1*Δ expressing *S. paradoxus* FKH2 (right), growing in rich conditions (YPD).

**Fig 4.**
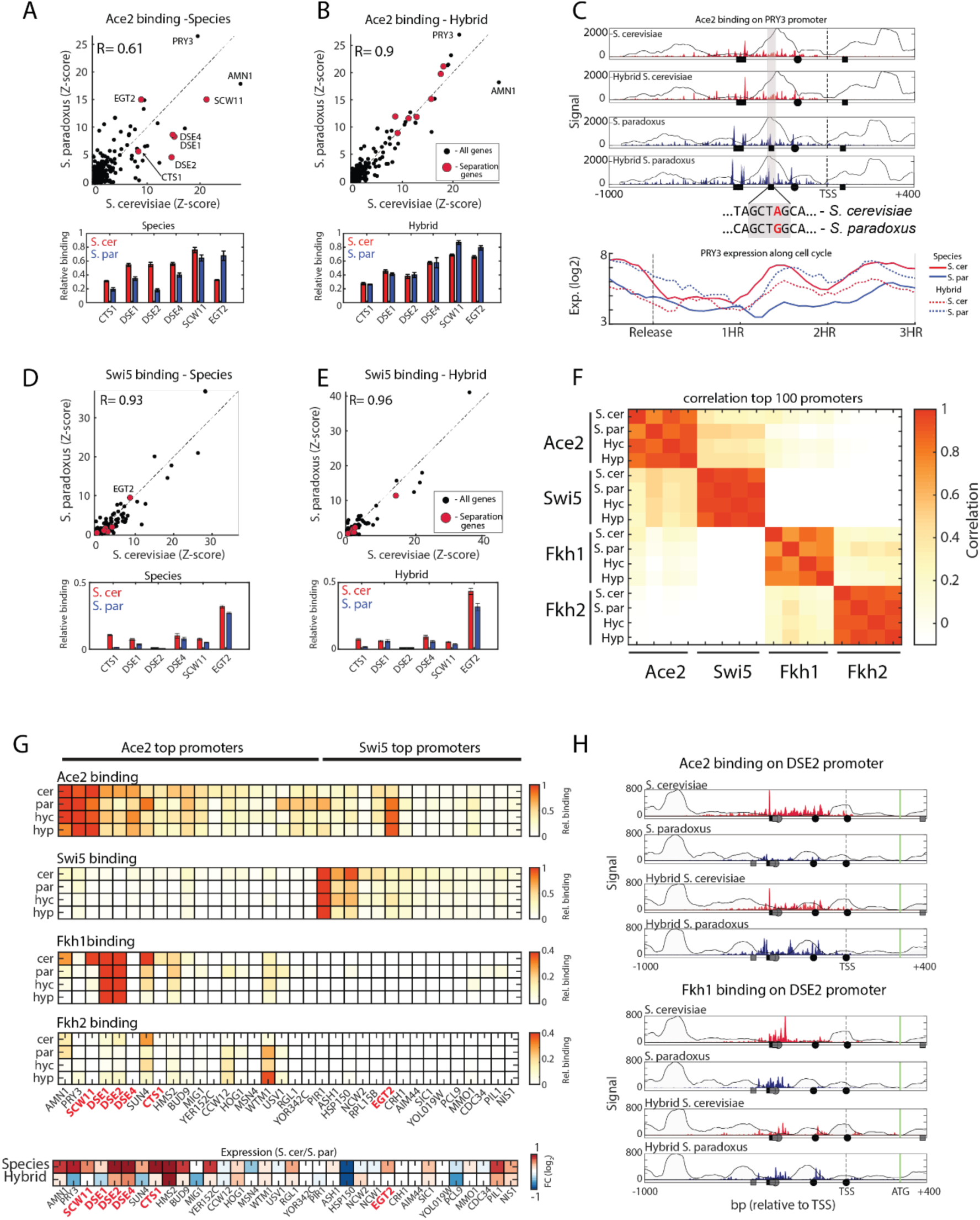
*Cis* and *trans* regulatory variations affect ACE2 binding. **(A)** ChEC-seq binding profiles of Ace2 between species. Top: plotted is Ace2 sum of signal on each promoter after Z-score normalization. Pearson correlation values of top 100 promoters are shown, genes involved in cell separation are marked in red. Bottom: Relative Ace2 binding on cell separation genes. **(B)** ChEC-seq binding profiles of Ace2 within the hybrid, Same as in (A). **(C)** Ace2 differentially binds PRY3 promoter. Top: Ace2 binding on PRY3 promoter, black circles and squares represent Ace2 binding motif (CCAGC) on + and – strand, respectively. Background line represents nucleosome occupancy. Dashed line represent transcription start site (TSS), x-ticks are bp relative to TSS. Middle: Sequence changes leading to loss or gain of Ace2 motif are shown. Bottom: PRY3 expression along the cell cycle, note the *cis* effect between hybrid alleles. **(D-E)** ChEC-seq binding profiles of Swi5 between species and within hybrid, same as in (A-B). **(F)** Pearson correlation matrix of top 100 promoters of Ace2, Swi5, Fkh1 and Fkh2. Hyc and Hyp refer to Hybrid *S. cerevisiae* genome and *S. paradoxus* genome, respectively. Note that paralogs show considerable correlations, Fkh1 show low positive correlation with *S. cerevisiae* Ace2. **(G)** Binding to top Ace2 and Swi5 promoters. Top: Shown are relative sum of signal on promoters for each factor in the species and hybrid. Note overlap of Fkh1 and Fkh2 with Ace2 promoters. Note that different color scale for Fkh1 and Fkh2 is shown. Bottom: Differences in expression levels. Note, cell separation genes (marked in red) show higher expression in *S. cerevisiae*, which is mostly lost in the hybrid (*trans* effect). **(H)** Binding of Ace2 and Fkh1 on the cell separation gene DSE2. Circles and squares represent Ace2 binding motif (CCAGC, black), and Fkh1 binding motif (GTAAACA, grey). Note *trans* effects both in Ace2 and in Fkh1.

**Fig 5.**
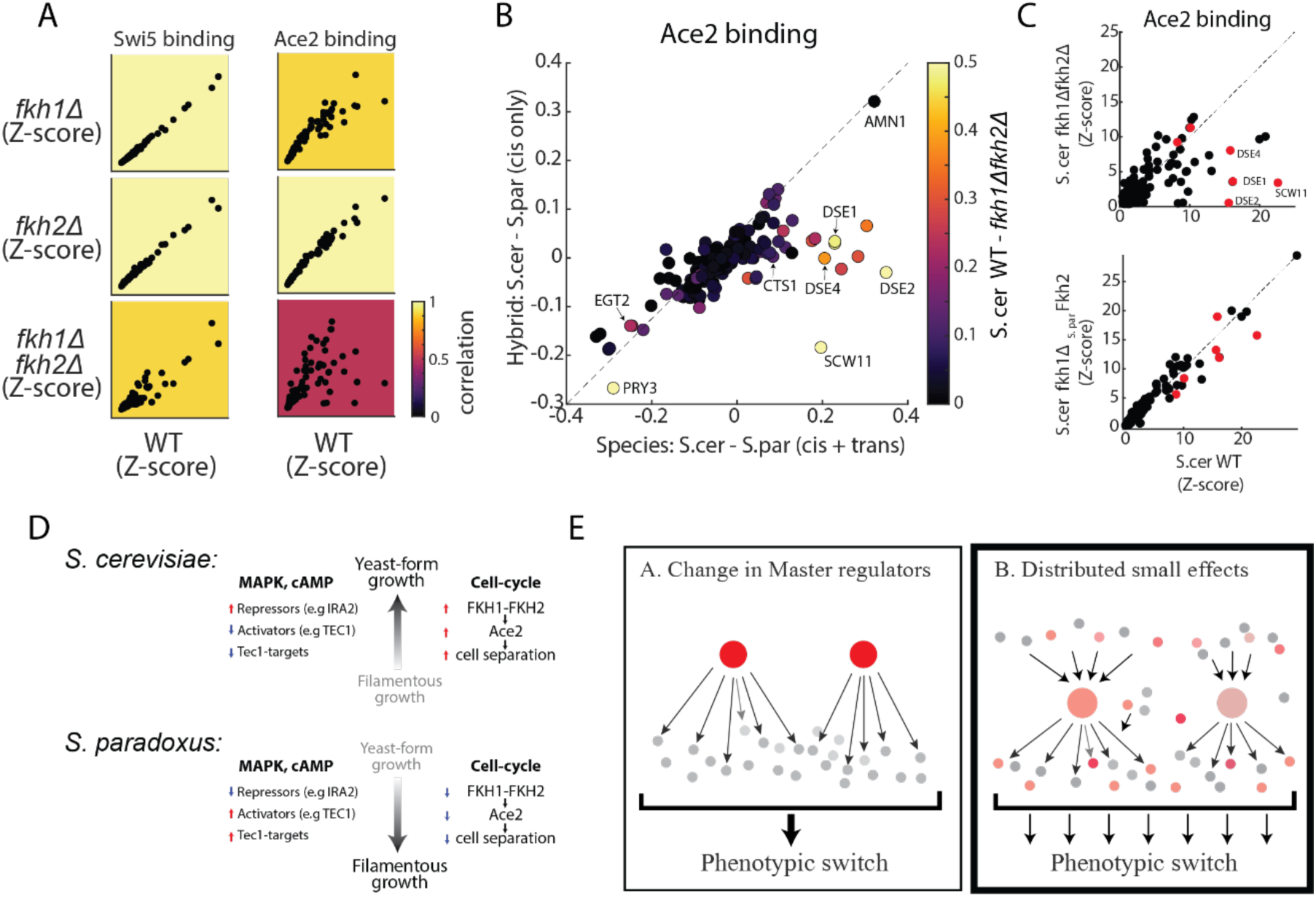
Fkh1 and Fkh2 directly mediates Ace2 binding to cell separation genes. **(A)** Ace2 binding is affected by Fkh1 and Fkh2. Shown are normalized sum of signal on promoters for Swi5 and Ace2 in WT *S. cerevisiae* compared to *fkh1*Δ, *fkh2*Δ and *fkh1*Δ*fkh2*Δ. Color represents correlation of top 100 promoters to WT. **(B)** *Cis* and *trans* effects on Ace2 binding. Scatter plot of relative Ace2 binding differences between species (x-axis) to within hybrid (y-axis). *Cis* effects are therefore along the diagonal while *trans* effects deviate from it. Color represents the differences in Ace2 binding between *S. cerevisiae* WT to *fkh1*Δ*fkh2*Δ. Note that cell separation genes show a strong *trans* effect and are strongly affected by *fkh1*Δ*fkh2*Δ. **(C)** Expression of *S. paradoxus* FKH2 in *S. cerevisiae* restores Ace2 binding. Scatter plots of normalized sum of signal on promoters of Ace2 in *S. cerevisiae* WT vs *fkh1*Δ*fkh2*Δ (top), and in *S. cerevisiae* WT vs. *fkh1*Δ expressing *S. paradoxus* FKH2 (right). Red dots represent cell separation genes. **(D)** overview of inter-species regulatory differences. Expression variations across different regulatory levels, affecting both filamentous regulation and cell cycle regulation, supporting the respected growth mode. Red and blue arrows indicate higher or lower expression, respectively.**(E)** Model of evolutionary adaption of complex phenotypes: While mutations that alter activity or expression of a master regulator could activate the cellular (A), we suggest that weak distributed effects accumulate in different layers of the regulatory network, driving and stabilizing the different response (B).

Overexpression of TEC1 in *S. cerevisiae* can induce filamentous growth, while its deletion prevents this transition even in nitrogen starved cells[23,43]. In order to see whether the changes between the species are seen also between *S. cerevisiae* in yeast or filamentous growth, we examined an expression dataset comparing *S. cerevisiae tec1Δ* to TEC1 overexpression[24]. Indeed, the majority of expression differences between the species was similar to changes between *S. cerevisiae* in yeast-form growth and filamentous growth (S2B Fig). We therefore hypothesized that the variations in the expression of Tec1-inducing MAPK pathway genes, and in TEC1 itself, may account for *S. paradoxus* filamentous phenotype. To examine that, we first asked whether Tec1 target genes (Defined from[24], Fig S2C) are expressed at higher levels in *S. paradoxus*. This was indeed the case, as these genes showed, on average, 1.25-fold higher expression in *S. paradoxus* (Fig 2D). We expected this expression difference to be lost in the hybrid, where Tec1 does not distinguish between alleles. This, however, was not the case, as the hybrid showed 1.2-fold higher expression from the *S. paradoxus* genome (Fig 2D), indicating on accumulated *cis* variations in Tec1-target genes. Further, deleting TEC1 did not reduce the expression of these genes in *S. paradoxus* (S2D Fig), indicating that *cis* expression variations in Tec1-targets were mostly independent of Tec1 itself. Consistent with that, TEC1-deleted cells showed no detectable effect on the filamentous phenotype (Fig 2E). Therefore, while variations in TEC1 correlate with *S. paradoxus* filamentous growth, Tec1 is not required for generating this phenotype.

The finding that Tec1 is dispensable for filamentous growth in *S. paradoxus* was surprising to us, as this TF is the central output of the MAPK cascade triggering filamentous growth in nutrient-starved *S. cerevisiae* cells. We therefore tested two additional TFs, Phd1 and Rim101, that control filamentous growth in *S. cerevisiae*, but are activated through pathways other than the MAPK cascade[44,45]. Neither PHD1, nor RIM101 were essential for filamentous growth in *S. paradoxus* (S2E Fig), though a mild reduction in flocculation was apparent. Therefore, key TFs which are essential for triggering filamentous growth in *S. cerevisiae* under poor conditions, are dispensable for this phenotype in *S. paradoxus* growing in rich conditions.

### *Cis*-effects bias expression of cell-cycle regulators

The finding that key upstream activators of the filamentous program are not required for maintaining this program in rich-media growing *S. paradoxus*, suggests that additional variations exist, acting downstream or in parallel to these signaling pathways. In *S. cerevisiae*, mutating cell cycle regulators can induce filamentous growth in a manner that is independent of Tec1[24,27,46]. Variations in cell-cycle regulators could therefore contribute to the yeast-filamentous divergence of the two species (Fig 3A). To examine this, we set out to compare the expression of cell-cycle regulators between the two species.

Cell cycle progression is driven by the periodic activation of the cyclin proteins[47]. We examined the expression patterns of the three G1 cyclins (CLN1-3) which act at the G1/S transition, and the six B-type cyclins (CLB1-6) acting in the later cell cycle phases (Fig 3B). Comparing expression levels revealed that six of the nine cyclins are differentially expressed between species. These inter-species differences in expression correlate with known cell-cycle phenotypes: the longer duration of the G2/M phase of *S. paradoxus* for example, corresponds to the reported consequences in *S. cerevisiae* of ectopically expressing CLB3 and CLB6, or deleting CLB1[48]. Similarly, CLN3, a principle driver of the G1/S transition, showed two-fold higher expression in *S. paradoxus*, consistent with its shorter G1. Examining the hybrid, we found that most variations are suppressed in this uniform *trans* environment (Fig 3B). Therefore, while cyclins expression correlates with the respective cell-cycle phenotype, these differences are not due to *cis* effects, but result from *trans* effects propagated from variations in upstream regulators.

We next considered the TFs regulating the periodic expression of cell-cycle genes (Fig 3C). Differential expression was observed for MBP1 (FC=2), which acts at the G1/S transition, and for FKH1 and FKH2 (FC=1.4, 2, respectively), which act at S/G2. All three TFs showed higher expression in *S. cerevisiae*. For MBP1 and FKH2, this difference in allele expression remained in the hybrid, pointing to *cis* variation. Testing reported targets of Fkh1 and Fkh2[46], we found that these are also more highly expressed in *S. cerevisiae* (average FC = 1.25), but were similarly expressed in the hybrid, as expected of *trans-*effects (Fig 3D, Fig S3A). Among these targets are the cell cycle paralogs TFs Ace2 and Swi5, which regulate the expression of late M-early G1 genes. Similar to other Fkh2 targets, expression of SWI5 and ACE2 was higher *S. cerevisiae* (FC = 1.75, 1.2, respectively), as was the expression of Ace2’s reported targets[49] (average FC= 2.6) but not Swi5’s (Fig 3D, Fig S2B). These effects were mostly lost in the hybrid, indicating their *trans* origin (Fig 3D). Together, these results suggest that *cis* variations in FKH2 might propagate to affect not only its own targets, but also targets of the downstream TF, Ace2.

The transcription cascade starting in Fkh1-Fkh2 and propagating through Ace2-Swi5 is central for the transition between yeast and filamentous growth[18]. In particular, Ace2 and Swi5 induce cytokinesis by expressing cell separation genes required for the full separation of mother and daughter cells during yeast-form growth[50,51]. Further, Ace2 inhibits the expression of CLN3 and by this extends the G1 duration of daughter cells, leading to the a daughter-specific delay characterizing yeast-form growth[49,52](Fig 3E). Indeed, deletion of ACE2 in *S. cerevisiae* reduced the portion of cells found in G1 and induces clump formation, as indicated by DNA staining (Fig 3F). Surprisingly, SWI5 deletion increased G1 duration in *S. paradoxus* and hybrid, but had no effect in *S. cerevisiae* (Fig 3F). Deletion of neither gene, however, lead to a switch in the growth-mode.

Fkh1 and Fkh2 contribute to yeast growth not only by inducing ACE2 and SWI5 expression, but also by activating expression of the CLB2-gene cluster, known to influence the transition between yeast and filamentous growth[53]. We therefore asked whether the lower expression of FKH1 and FKH2 in *S. paradoxus* accounts for its filamentous phenotype. *S. cerevisiae* cells deleted of both Fkh1 and Fkh2 became filamentous even in rich conditions, consistent with previous reports[46,53] (Fig 3G). However, expressing FKH2 using the *S. paradoxus* promoter was sufficient for retrieving yeast growth to the FKH1-FKH2 deleted cells (Fig 3G), although FKH2 expression levels were 2-folds lower then when expressed using the *S. cerevisiae* promoter (S3C Fig). Therefore, the low FKH1 and FKH2 expression cannot, by itself, account for *S. paradoxus* filamentous phenotype.

### *Cis*-effects bias expression of transcription-factor cascade regulating cell-separation

Low-level expression of FKH2 is therefore sufficient for retrieving yeast growth for *S. cerevisiae*, but not for *S. paradoxus*. It was previously reported that Fkh1 and Fkh2 bind to Ace2 target promoters and function as Swi5-specific anti-activators[54]. We therefore hypothesized that the lower expression levels of FKH1 and FKH2 may alter the activity of Swi5 and Ace2. As a measure of this activity, we measured the TFs binding profiles via ChEC-seq[55] in both species and hybrid, and examined whether binding specificities are conserved. We began this analysis with Ace2, considering its direct role in regulating cell separation after cytokinesis.

Binding profiles highly correlated between repeats (R= 0.94-0.97 on average), and were localized upstream to the transcription start site (TSS; S4A and S4B Figs). While the preferred Ace2 binding motif in both species was identical to the known Ace2 motif (S4C Fig; CCAGC[56]), the overall promoter selection has changed (Fig 4A top; promoter selection correlation=0.61). More specifically, promoters of cell-separation genes (CTS1, DSE1, DSE2, DSE4 and SCW11, reviewed in[50]) were bound by Ace2 more strongly in *S. cerevisiae* than in *S. paradoxus* (Fig 4A bottom; 1.8-fold higher signal on average). Notably, EGT2, the only Swi5-regualted cell separation gene[50], did not conform to this general behavior and showed higher binding signal in *S. paradoxus*. Therefore, the reduced expression of cell separation genes in *S. paradoxus* is not only due to the lower ACE2 levels, but also due to changes in its binding specificity between the two species.

We hypothesize that the higher expression level of the ACE2 in *S. cerevisiae* may affect its binding specificity. To test this, we first reduced ACE2 expression in a diploid *S. cerevisiae* by deleting one of its alleles, reaching levels even lower than observed in *S. paradoxus*. This, however, had little effect on its binding specificity (S5A Fig; R=0.94). Similarly, within the hybrid, the two Ace2 orthologs were localized to the precise same locations (S5B Fig; R = 0.99). Therefore, differences in Ace2 binding specificity in the two species are not due to differences in Ace2 expression level or protein sequence.

Binding specificity of Ace2 diverged significantly between species (R=0.61, Fig 4A). We therefore asked whether these variations result from *cis* mutations, or from variations in a *trans* acting factor. To examine this, we compared Ace2 binding to the two genomes within the hybrid (Fig 4B top). The majority of differences in Ace2 promoter binding specificities were suppressed within the hybrid, indicating on *trans* acting variation (Fig 4B; R =0.9). In particular, cell separation alleles of both *S. cerevisiae* and *S. paradoxus* were bound by Ace2 to a similar extent (Fig 4B bottom). *cis* effects were observed in a small number of genes. PRY3, for example, a cell wall associated protein, was more strongly bound (1.4-fold) in *S. paradoxus*, while AMN1, a modulator of cell separation and mitotic exit, showed preferred binding (1.5-fold) to the *S. cerevisiae* promoter (Fig 4C and S5C). In both cases, one Ace2 binding site was absent from the promoter showing lower binding. Further, expression of the respective alleles varied in *cis* (within the hybrid), supporting the functionality of this gain (or loss) of binding site. We also noted reduced Ace2 binding to the CLN3 promoter, which was associated with the loss of a well-studied Ace2 binding site[49] (S5D Fig). This binding site was previously implicated in extending G1 duration, suggesting that its loss in *S. paradoxus* may account for its short G1. Mutating this binding site to match the *S. paradoxus* allele lead to a mild but significant reduction (5%) in G1 duration in *S. cerevisiae* as measured both by DNA staining and live microscopy (S5D-F Figs). Yet this cannot account, by itself, for the reduced G1 duration of *S. paradoxus* (50% reduction compared to *S. cerevisiae*[20]).

The lower binding of Ace2 to cell-separation promoters in *S. paradoxus* is therefore due to variations in *trans*. We next asked whether such *trans* variations affect also the binding of Ace2’s paralog, Swi5. Swi5 was bound to the same DNA motif as Ace2 in both species (S4C Fig). In contrary to Ace2, Swi5 promoters binding specificities were largely conserved between the species and within the hybrid (R=0.93,0.96, respectively, Figs 4D and 4E top). While showing a considerable overlap with Ace2 promoters (Fig 4F), Swi5 binding to cell separation promoters was significantly weaker than that of Ace2 (Figs 4D and 4E bottom). Therefore, the *trans* variations affect Ace2 binding specifically, without affecting its paralog, Swi5.

### Fkh1 and Fkh2 mediate Ace2 binding to cell-separation promoters

Fkh1 and Fkh2 were previously shown to bind Ace2-specific promoters and block their activation by Swi5[54]. We therefore hypothesized that Fkh1 and Fkh2 may regulate Ace2 activity by promoting its binding to specific promoters, including cell-separation promoters. Profiling the binding of Fkh1 and Fkh2 revealed that while these TFs bind a distinct set of promoters than Ace2 and Swi5 (Fig 4F), cell-separation genes were bound by Fkh1, and to lesser extent by Fkh2 (Figs 4G and 4H). Additionally, these genes showed higher expression in *S. cerevisiae* than in *S. paradoxus*, which was mostly buffered in the hybrid (Fig 4G bottom). Overall, cell separation promoters that are more strongly bound by Ace2 in *S. cerevisiae*, are bound also by Fkh1 and FKH2 but not by Swi5 in both species and in the hybrid.

To examine whether Fkh1 and Fkh2 recruit Ace2 to cell-separation promoters, we examined *S. cerevisiae* cells deleted of FKH1, FKH2 or both. Deleting either FKH1 or FKH2 had little effect on binding of Ace2 and Swi5 (R = 0.92-0.99, Fig 5A). Swi5 binding was also largely invariant to the deletion of both factors (R = 0.93), yet Ace2 binding to its top promoters was largely suppressed (R = 0.53, Fig 5A). More specifically, deletion of both FKH1 and FKH2 affected Ace2 binding primarily in promoters showing high *trans* variations (Fig 5B). Since these double-deletion cells are filamentous, we controlled for non-specific effects of the filamentous phenotype by over-expression of CLB2 in this background[53]. While this overexpression retrieved yeast growth, Ace2 did not regain binding to cell-separation promoters (S6B and S6C Figs). Together, these results suggest that Fkh1 and Fkh2 directly regulate the recruitment of Ace2 to promoters, and play a role in the inter-species *trans* variations of Ace2 binding.

We reasoned that the low levels of FKH1 and FKH2 in *S. paradoxus* could explain the reduced binding of Ace2 to cell-separation promoters in this species. Contrasting this possibility, however, low FKH2 levels expressed using the *S. paradoxus* promoter in *S. cerevisiae* were sufficient to retrieve Ace2 binding to FKH1/2-deleted cells (Figs 5C and S3C). Therefore, an additional layer of variations exists, whereby low levels of FKH2 are sufficient to promote Ace2 binding in *S. cerevisiae* but not sufficient for its recruitment in *S. paradoxus*. These variations could act in *trans* through additional signaling or other unknown factor, or could act in *cis* by modulating Ace2-Fkh1/2 promoter interactions. Further experiments are required to test that.

## Discussion

A major challenge in the study of regulatory evolution is to link variation in gene expression to differences in phenotypes. Here, we examine this by comparing budding yeast species that express largely the same set of genes and exhibit similar growth requirements, yet show major differences in cell-cycle dynamics, budding pattern, cell morphology, and adhesion properties when growing in the same environment. While these differences between the species encompass a wide range of cellular functions, they can be collectively explained as two distinct growth modes, yeast and filamentous, available to the individual species. This suggests that this major divergence results from the differential activation of a pre-existing differentiation program, rather than optimization of each property individually. Regulators of the yeast-to-filamentous transition are well characterized, providing a tractable model for analyzing systematically the relation between variation in gene expression and differences in phenotypes.

The transition to filamentous growth is regulated by evolutionarily conserved signaling pathways[17,57]. One of these is the MAPK cascade that activates the transcription factor, Tec1. We found that most activators in this pathway are expressed at higher levels in *S. paradoxus*, while inhibitors expression is higher in *S. cerevisiae* (Fig 5D). Variants of one such inhibitor, Ira2, was previously reported to be involved in morphological variation in wild *S. paradoxus* isolates [21], supporting the notion that these expression variations could have a large phenotypic effect. Expression of TEC1 itself was 4-folds higher in *S. paradoxus*, and this was largely due to gene-linked mutation (*cis* effect). This, together with the ability to induce filamentous growth through TEC1 overexpression, poised Tec1 as a promising candidate for driving the phenotypic difference. However, contrasting our expectation, Tec1 is in fact dispensable for *S. paradoxus* filamentous growth. Examining Tec1-target genes, as defined in *S. cerevisiae*, provided a partial explanation for this result, as these genes accumulated *cis* mutations rendering their higher expression in *S. paradoxus* largely independently of Tec1. Therefore, while genes of the MAPK and its targets accumulated *cis* mutations that biased their expression in the direction favoring the respective phenotype, its downstream-acting TF is in fact not sufficient for explaining the difference in phenotype.

Subsequent analysis of cell-cycle regulators led to a similar conclusion; we noted that expression of the TFs FKH1 and FKH2 is lower in *S. paradoxus* and this reduced expression propagates downstream, affecting their targets, as well as targets of their downstream TFs, Ace2 and Swi5 (Fig 5D). Furthermore, we found that Fkh1 and Fkh2 not only activate ACE2 and SWI5 expression, but also recruit Ace2 to cell separation genes specifically in *S. cerevisiae*. Yet, while *S. cerevisiae* cells deleted of both FKH1 and FKH2 are filamentous, expressing FKH2 using the weak *S. paradoxus* promoter was sufficient to retrieve yeast growth.

Based on these results, we propose that the initial steps of this differential recruitment did occur through a genetic shortcut whereby a single regulator was modified to activate or repress the phenotypic switch in a new context, as often seen in experimental evolution assays[58,59]. However, once this program has been initiated, additional mutations continued to accumulate and spread throughout the regulatory network. This way, variations that bias expression in the ‘wrong’ direction will be eliminated by purifying selection, while those which promote the selected program will have a higher chance to persist, and will compensate for possible deleterious mutations in the initial driver. Indeed complex traits are mostly polygenic, and adaptive selection on such a trait is predicted to affect many loci[12,15]. The resulting distributed stabilization may explain the difficulty in identifying causal variations, and the prominence of both *cis* and *trans* variations that correlate with the phenotype[60].

Our model, proposing distributed stabilization of adapted complex traits, follows the conceptual framework of the omnigenic model for genetic variations[13,61] (Fig 5E). The omnigenic model explains the abundance of weak effects in genetic association studies as a consequence of the high connectivity within the transcriptional network. This connectivity expands the space of variations propagating to modulate the principle phenotypic-relevant pathway. In this framework, each individual carries a different representative of this possible pool of variations, explaining the small effect of each variant. In our case, we consider variations that have all been stabilized within the same species. The abundance of variations whose effect can influence the respective program results in a compensatory manner, which spreads the effect of each individual mutation. Therefore, each individual variation will have a small effect, leading to a distributed stabilization of all effects. Further studies are required to examine whether this model applies also in commonly studied cases such as evolutionary co-option that shape the body pattern.

## Materials and Methods

### Strains and plasmids

All strains used in this study and their genotypes are listed in Table S1. For timecourse experiments, Strains used were *S. cerevisiae* of background BY4743 (Diploid) and *S. paradoxus* of background CBS432 (Diploid). Hybrid was created by mating *S. cerevisiae* Mat a with *S. paradoxus* Mat alpha.

For ChEC-seq experiments, each transcription factor (TF) was C-terminally tagged with MNase. Yeast cells were transformed with the amplification product of MNase-Kanamycin cassette from pGZ108 plasmid, a gift from Steven Henikoff. Standard transformation using 50bp homology-based recombination was used. Hybrids were created by mating haploid TF-MNase strain, both of *S. cerevisiae* and *S. paradoxus*, with WT haploids of the other species to create reciprocal hybrids, each expressing a TF-MNase allele of either species. ChEC-seq experiments on Ace2, Swi5, Fkh1 and Fkh2 were done on haploid strains. Ace2 and Swi5 were additionally checked in diploid strains (expressing one copy of TF-MNase) to control for ploidy effects.

Gene deletions were generated by replacing each relevant gene’s ORF by transformation and growth on plates containing selection. Strains of genotype *fkh1Δ*::hph and *fkh2Δ*::hph were generated by amplifying the hph gene from plasmid pBS35. Double deletion of genotype *fkh1Δfkh2Δ* were generated by amplifying the LEU2 gene from plasmid pRS425 and replacing FKH2’s ORF. Strains of genotype *ace2Δ*::KanMX and *swi5Δ*::KanMX were generated as reported at Krieger et al[62].

Swapped FKH2 strain was created in two steps: first, the FKH2’s ORF of *S. cerevisiae*, including terminator and promoter (150bp downstream and 700bp upstream, respectively), was replaced by an amplicon of the KanMX gene from plasmid pBS7. Second, a CRISPR based method was used to induced break in the KanMX gene, while co-transforming with a genomic amplicon of FKH2 (including terminator and promoter) from *S. paradoxus*. Cas9 and the locus-specific 20bp gRNA were expressed using plasmid bRA89. Ligation of the gene-specific gRNA into the bRA89 plasmid was done as previously described[63].

All strains were validated by PCR and sequencing.

### Microscopy

Yeast cells were grown in YPD over night at 30°C to stationary phase, and were inoculated to fresh medium for a few hours until reaching OD600 of 0.2. The cells were then prepared for imaging on YPD 2% low-melt agar pads in 96-well plate. Images were taken either in an Olympus IX83 based Live-Imaging system equipped with CSU-W1 spinning disc: sCMOS digital Scientific Grade Camera 4.2 MPixel Growth or Zeiss Axio Observer Z1 inverted microscope equipped with a motorized XY and Z stage, external excitation and emission filter wheels (Prior), IR-based Definite Autofocus from Zeiss and a 63 × oil objective. The cells were kept at 30 °C. Images were taken every 5 minutes. image adjustments and labeling were performed using imageJ[64].

### Time-course experiments

Yeast cells were grown in YPD over night at 30°C to stationary phase, and were inoculated to fresh medium to OD600 of ∼0.005. When reaching an OD600 of 0.1-0.2, hydroxyurea (HU) was added to the media to a final concentration of 0.2M for additional 2 hours. To remove HU from the media, the cells were washed twice from by centrifugation (4000rpm for 1 minute) and re-suspended in fresh, warm, equal-volume YPD. Then, the culture was returned to a bath orbital shaker. Cells were collected at the following time points: before HU, 5’,10’,20’,30’,60’ and 120 minutes in HU, and every 5 minutes after release for 3 hrs. In total 43 time points for each strain. For RNA, samples of 1.5ml were taken and centrifuged for 10 seconds in 13,000 rpm, sup was removed and the pellets were immediately frozen in liquid nitrogen. For DNA staining, samples of 1.5ml were taken and centrifuged for 10 seconds in 13,000 rpm and resuspended in cold 70% ethanol and kept in 4°C. This experiment was carried with two independent biological repeats for each strain.

### Flow Cytometry – DNA staining

Cells were washed twice with 50mM Tris-HCl pH8, re-suspended in RNase A for 40 minutes in 37°C, washed twice with 50mM Tris-HCl pH8, and re-suspended in Proteinase K for 1-hour incubation at 37°C. Then, cells were washed twice again, and re-suspended in SYBR green (1:1000) and incubated in the dark at room temperature for 1 hour. Next, cells were washed from the stain, re-suspended in 50mM Tris-HCl pH8 and sonicated in Diagenode bioruptor for 3 cycles of 10’’ ON and 20’’ OFF in low intensity. Cells were taken to FACS for analysis using BD LSRII system.

### RNA extraction and sequencing

RNA was extracted using a modified protocol of nucleospin^®^ 96 RNA kit (Machery-Nagel, cat 740466.4). Specifically, cells lysis was done in a 96 deep-well plate by adding 450µl of lysis buffer containing 1M sorbitol (SIGMA-ALDRICH), 100mM EDTA and 0.45µl lyticase (10IU/µl). The plate was incubated in 30°C for 30 minutes in order to break the cell wall, centrifuged for 10’ at 2500 rpm, and supernatant was removed. From this stage, extraction proceeded as in the protocol of nucleospin^®^ 96 RNA kit, only substituting β-mercaptoethanol with DTT. cDNA was prepared from the RNA extracts, barcoded, and sequenced using either Illumina HiSeq 2500 or Illumina NextSeq 500.

### Processing and analysis of RNAseq data

A pipeline for RNAseq data was created by Gil Hornung (INCPM, WIS). Fastq files were pre-processed by Merging them into one file, removal of reads with high A or T content, trimming of the first 3 bases (usually GGG) and trimming of Illumina adapters and bases with quality <10 using cutadapt.

The reads were then mapped against a dual-species reference genome of *S. cerevisiae* and *S. paradoxus* strain CBS432[65]. Genomes fasta and annotation files were downloaded from http://sss.genetics.wisc.edu/cgi-bin/s3.cgi. Mapping was performed with STAR 2.4.2a with the parameters --sjdbOverhang 60 --scoreGap -10. The alignments were divided based on the alignment scores. If for a read the highest scoring alignment is assigned to a certain genome, and is unique in that genome, then it is assigned to that genome. If there is no difference in the scores between the two genomes and the alignment is unique in the cerevisiae genome, then the alignment to the cerevisiae genome is kept and save as “indistinguishable”. On average, 85% of aligned reads were mapped to either genome. Indistinguishable reads were discarded from further analysis.

Counting was performed on the TES (Transcript End Site) region of each gene. The TES region is defined as 200 bp downstream from the end of the gene, and 500 bases upstream to the end of the gene. The reads were counted using htseq-count, with parameters --stranded yes and --mode union. The sum of all reads in each sample was normalized to be 1,000,000 and genes with expression below a threshold of log2(10) were excluded. All further data analysis was performed in Matlab.

### Expression of periodic genes

Periodic transcripts were defined by Cyclebase[39], and classified to groups based on Cyclebase expression peak time score. The top 500 genes were taken and intersected with genes that were detectable in our experiment for both species, resulting in ∼430 periodic genes. For determining if a gene show periodic expression, these 430 gene were classified into 12 groups based on expression peak time, the average expression along the timecourse of each group was calculated and the pearson correlation between each gene to the average of its group was calculated. Significant correlations were those with corrected p-value < 0.05 (after Benjamini Hochberg FDR procedure), and rho >0.4.

### Expression of phenotype associated genes

Genes with positive and negative effect of filamentous growth were assembled from the Saccharomyces Genome Database[42] (SGD). Positive regulators were defined as genes that when deleted, lead to reduced filamentous growth, or when over-expressed lead to an increase in filamentous growth. Negative regulators were defined as genes that when deleted, lead to an increase in filamentous growth, or when over-expressed lead to reduced filamentous growth. Only genes that were listed under ‘experiment type: classical genetics’ were used.

TEC1 regulated genes were computed from Madhani et al.[24] by comparing TEC1-over expression strain to that of *tec1Δ*. Genes above noise level were taken and sorted by their fold change difference (Fig S2A).

### Gene expression of mutants

Gene expression of *S. paradoxus* WT, *tec1Δ, phd1Δ* and *rim101Δ*, was taken from Krieger et al[62], using the average expression for two biological repeats for each mutant during exponential growth in YPD.

### Differential expression analysis

Differential expression analysis was carried out using DESeq2[66] on R 3.6.3. Read counts were given as input to DESeq2 with the design: “∼ genotype”, where “genotype” differentiates the sample as coming from: cerevisiae, paradoxus, hybrid-cerevisiae, hybrid-paradoxus. This design enabled differential expression analysis between species and between hybrid alleles (cis effect). Differential expression was determined via likelihood ratio test (test=“LRT”) focusing only on the interaction term For both designs, results went through log2 fold change shrinkage using ashr method[67].

### ChEC-Seq experiments

The experiments were performed as described previously[55], with some modifications. Cultures were grown overnight to saturation in YPD media and diluted into 50 ml of fresh YPD media to reach OD_600_ of ∼0.3 the following morning after ∼10 divisions. Cultures were pelleted at 1500 g and resuspended in 1 ml Buffer A (15 mM Tris pH 7.5, 80 mM KCl, 0.1 mM EGTA, 0.2 mM spermine, 0.5 mM spermidine, 1 × Roche cOmplete EDTA-free mini protease inhibitors, 1 mM PMSF), and then transferred to DNA low-bind tubes (Eppendorf 022431021). Cells were washed twice more in 500 μl Buffer A, pelleted, and resuspended in 200 μl Buffer A containing 0.1% digitonin. Then, cells were transferred to an Eppendorf 96-well plate (Eppendorf 951020401) for permeabilization (30 °C for 5 min). CaCl_2_ was added to a final concentration of 2 mM for 30 seconds for MNase activation. Next, 100 μl of stop buffer (400 mM NaCl, 20 mM EDTA, 4 mM EGTA and 1% SDS) were mixed with 100 μl sample. Proteinase K (100 μg, Sigma P2308) was then added, and incubated at 55°C for 30 min. Nucleic acid extraction was performed as previously described[55], with some modifications in the ethanol precipitation step; Samples were precipitated (at -80 °C for >1 hour) with 2.5 volumes of cold EtOH 96%, 45 μg Glycoblue (Thermo Fisher AM9515) and sodium acetate to a final concentration of 20 mM. DNA was centrifuged, washed with ethanol and treated with RNase A (Sigma, R4875) as previously described, followed by another round of DNA cleanup and ethanol precipitation. In order to enrich for small DNA fragments, reverse 0.8X SPRI clean-up was carried out. Library preparation was done as previously reported[68], except for the clean-up steps, which were performed using phenol-chloroform followed by ethanol precipitation as described above (instead of S400 columns). 1X SPRI was carried out on ChEC amplified libraries, which were then pooled and cleaned from adaptors dimers (150bp) if needed. Libraries were sequenced on Illumina NextSeq500 for paired end sequencing (50 bps for read1 and 15 or 25 bps for read 2).

Library construction for of ChEC-seq of Hybrid strain of FKH1-MNase and FKH2-MNase was slightly modified and phenol-chloroform clean-ups were replaced by SPRI-isopropanol clean-ups[69]. All ChEC-seq experiments were repeated at least twice for each strain.

### ChEC-Seq processing and analysis

Reads were aligned using Bowtie2[70] (parameters: –best –m 1) to a dual-species reference genome of S. cerevisiae R64 and S. paradoxus CBS432[71]. ChEC-Seq tracks, representing the enrichment of each TF, were calculated by adding +1 to each genomic location corresponding to the first nucleotide in a forward read, or the 50th position corresponding a reverse read. The signal was normalized to a total of 10 million reads, to control for sequencing depth. The median signal across repeats for each strain was taken for further analysis. For promoter analysis, promoters were defined only for genes with an annotated transcription start site (TSS)[72]. The length of each promoter was defined as 700 bps upstream to the transcription start site (TSS). The signal across each promoter was summed to calculate overall promoter binding for each sample.

## Acknowledgments

We thank all memebers of the Barkai lab. We thank Gil Hornung (INCPM, Weizmann Institute of Science) for constructing the RNA-seq pipeline. We thank Sagie Brodsky for both technical and computational help in ChEC-seq experiments and data analysis. We thank Dana Bar-Zvi, Tamar Jana, Gilad Yaakov and Miri Carmi for additional technical help and fruitful discussions.

## Supplementary figures

**S1 Fig.**
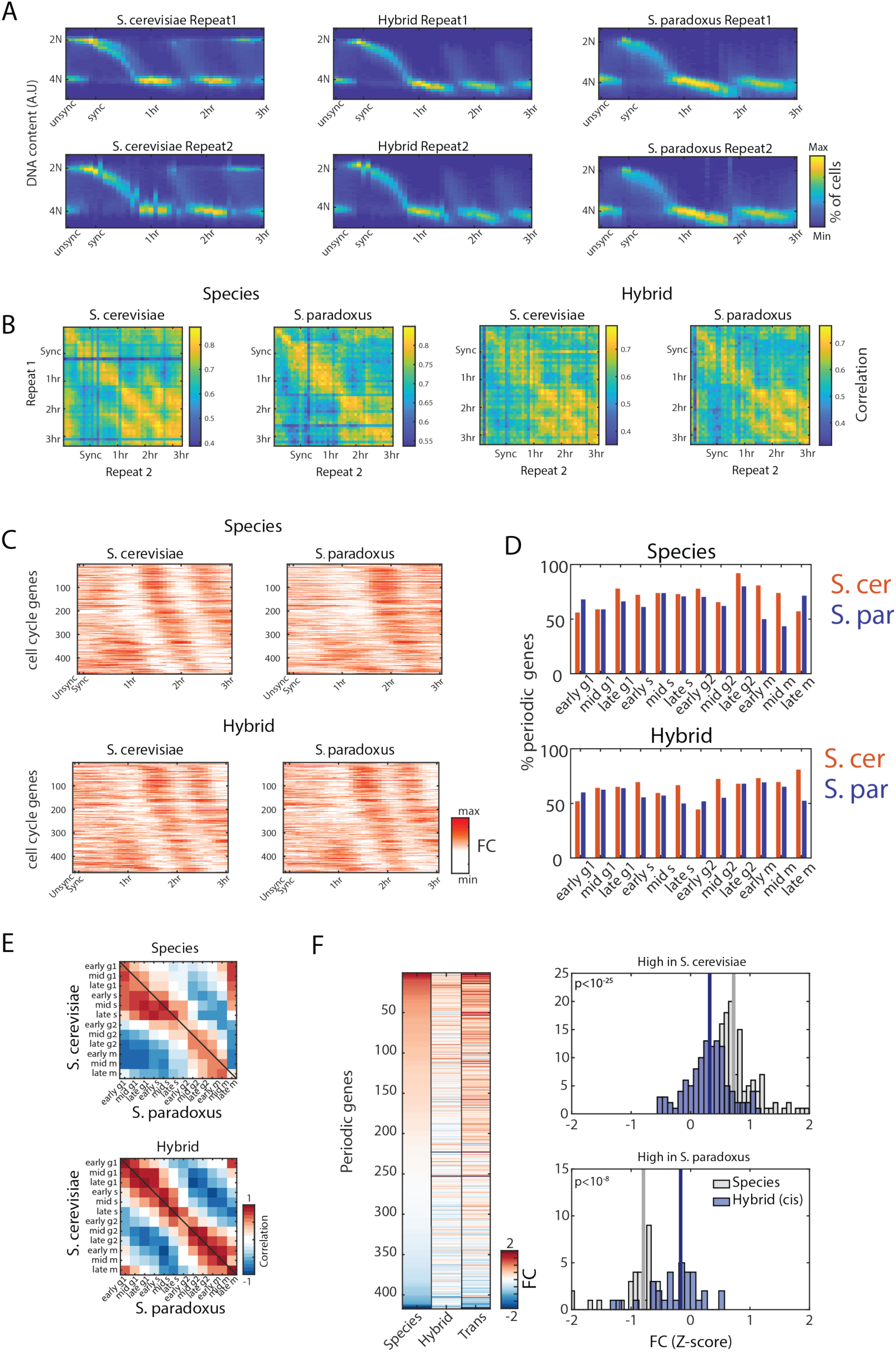
Monitoring the gene expression program along the cell cycle. **(A)** DNA staining profiles along the cell cycle in the two repeats of *S. cerevisiae, S. paradoxus* and the hybrid**. (B)** Correlation matrices of periodic transcripts (top 500) between the two repeats. Note high correlation along the diagonal and high correlation between first and second cycle. **(C)** Expression of periodic transcripts. Each gene was normalized to its median expression, plotted are all periodic genes ordered by their expression peak time. **(D)** Bar graphs showing the percentage of genes with significant correlation to the respected module (corrected p-value < 0.05, rho>0.4). **(E)** Periodic co-expression of cycling genes. Interspecies (left) and within hybrid (right) correlations between mean expression of cell cycle modules, classified to 12 groups. Note high correlations between early G1 to late genes in *S. paradoxus*, indicating rapid activation of S-phase genes. **(F)** Fold changes in expression levels of periodic genes. Left: shown are fold change (FC) in absolute levels at expression peak between species, hybrid and *trans* (Species-hybrid). Genes ordered by high expression differences in *S. cerevisiae*. Right: distribution of top changing genes in *S. cerevisiae* (top) and *S. paradoxus* (bottom).

**S2 Fig.**
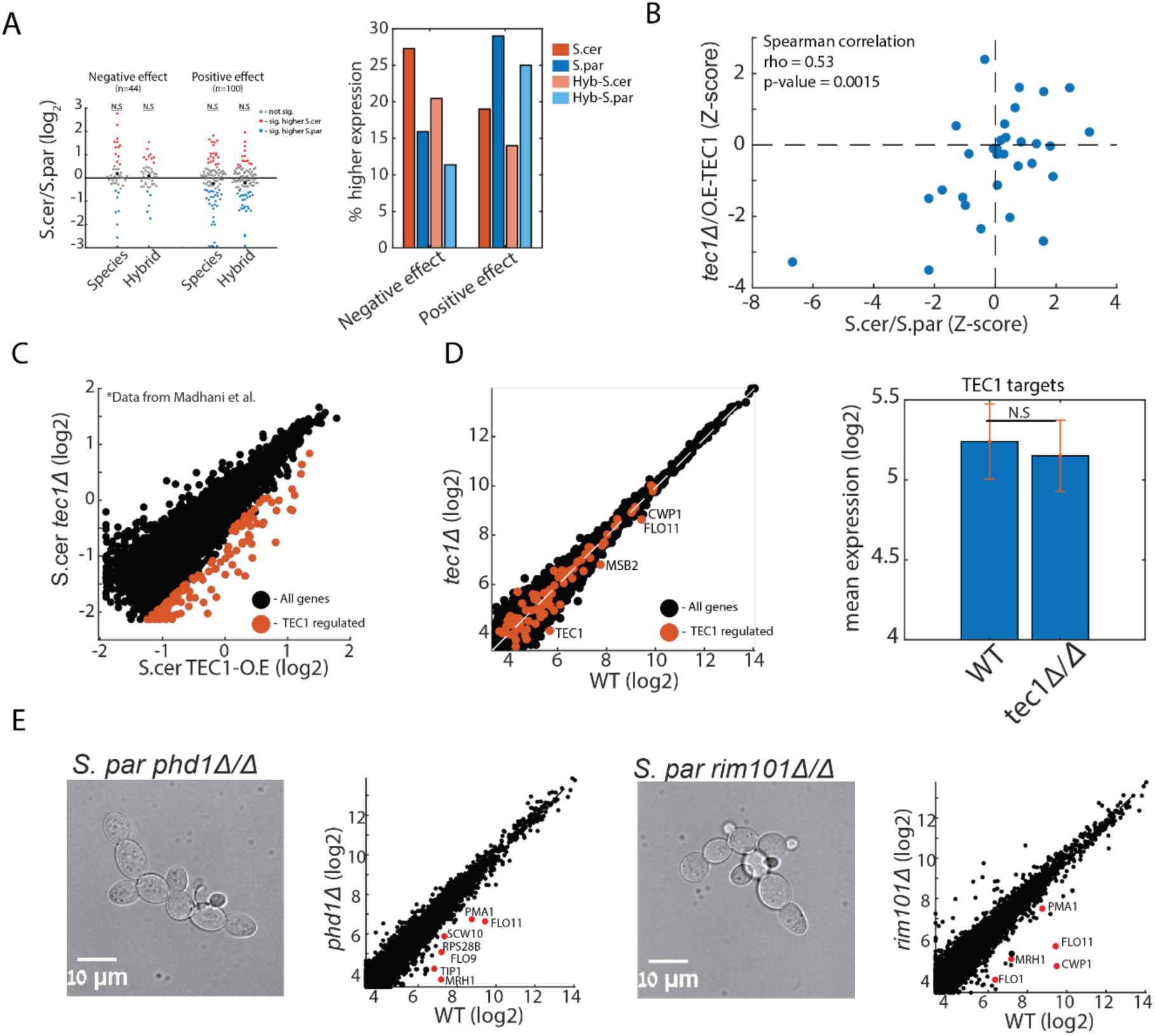
Expression of filamentous regulators. **(A)** Expression variations in genes that have reported negative (n=44) or positive (n=100) effect on filamentous growth. Left: Shown are average fold-change (FC) distributions across all timepoints. Red and blue dots indicate significantly higher expression in *S. cerevisiae* and *S. paradoxus*, respectively. Right: percentage of differential expressed genes in each group, note bias in *cis.* **(B)** Expression variations between species is similar to expression changes between *S. cerevisiae* in yeast-growth or filamentous growth. Shown are FC values of genes in MAPK and cAMP-PKA pathway between species (x-axis) to between *tec1*Δ to TEC1-over expression (y-axis). Data is presented in Z-scores. **(C)** Determining Tec1-regulated genes. Genes were defined as the genes showing significant change between TEC1-over expression to *tec1*Δ. **(D)** Expression of Tec1-regulated genes in *S. paradoxus* WT and in *tec1*Δ/Δ. **(E)** Phenotype and expression of *rim101*Δ/Δ and *phd1Δ/*Δ in *S. paradoxus* grown in YPD. Cells still exhibit filamentous growth, while cell-wall genes (such as FLO11, FLO9, CWP1) show lower expression.

**S3 Fig.**
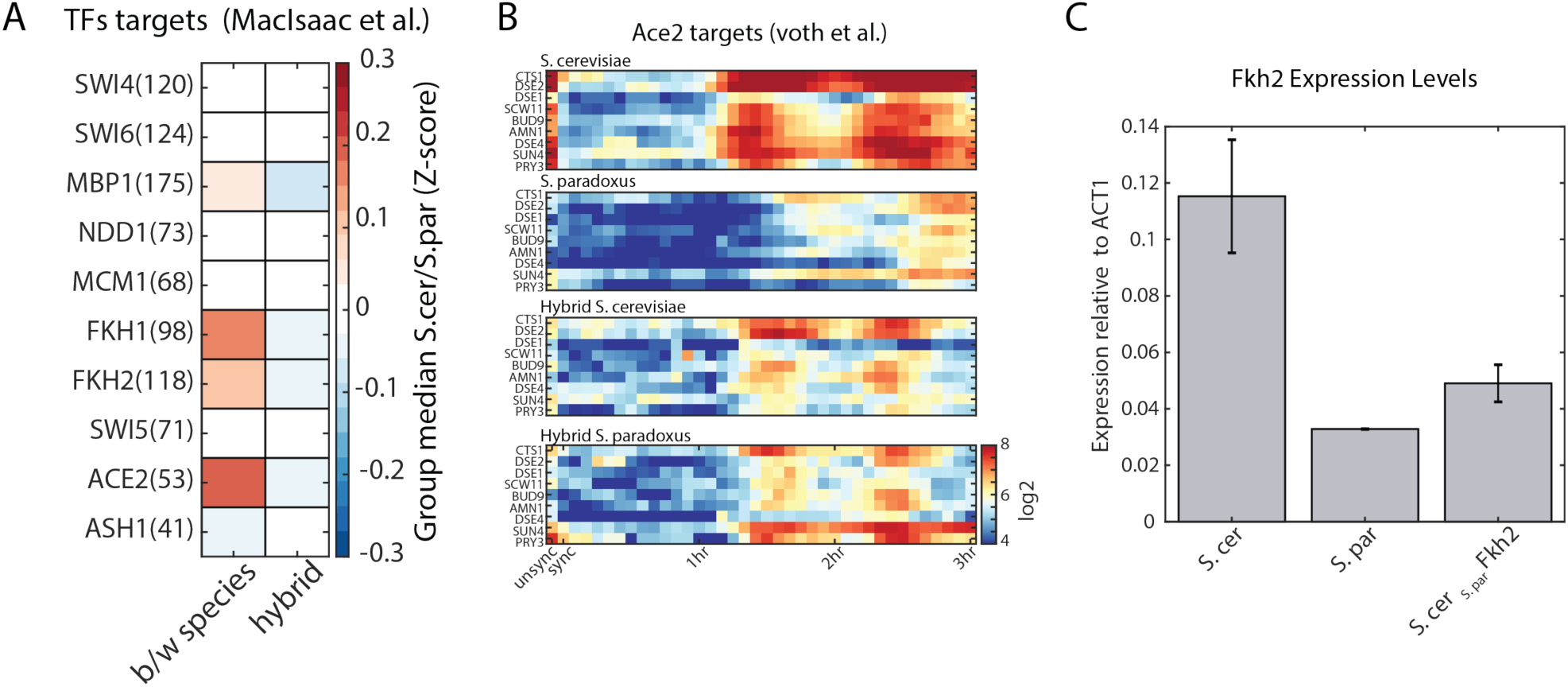
Downregulation of ACE2 targets in S. paradoxus. **(A)** Median FC (indicated in Z-scores) of TFs targets defined by another dataset ([73]). Number indicates the number of targets genes in the respected group. **(B)** Absolute expression levels of Ace2 targets (as defined by [54]) along the cell cycle (**C)** FKH2 expression levels, as measured by RT-qPCR, *in S. cerevisiae, S. paradoxus* and *S. cerevisiae* expressing *S. paradoxus* FKH2.

**S4 Fig.**
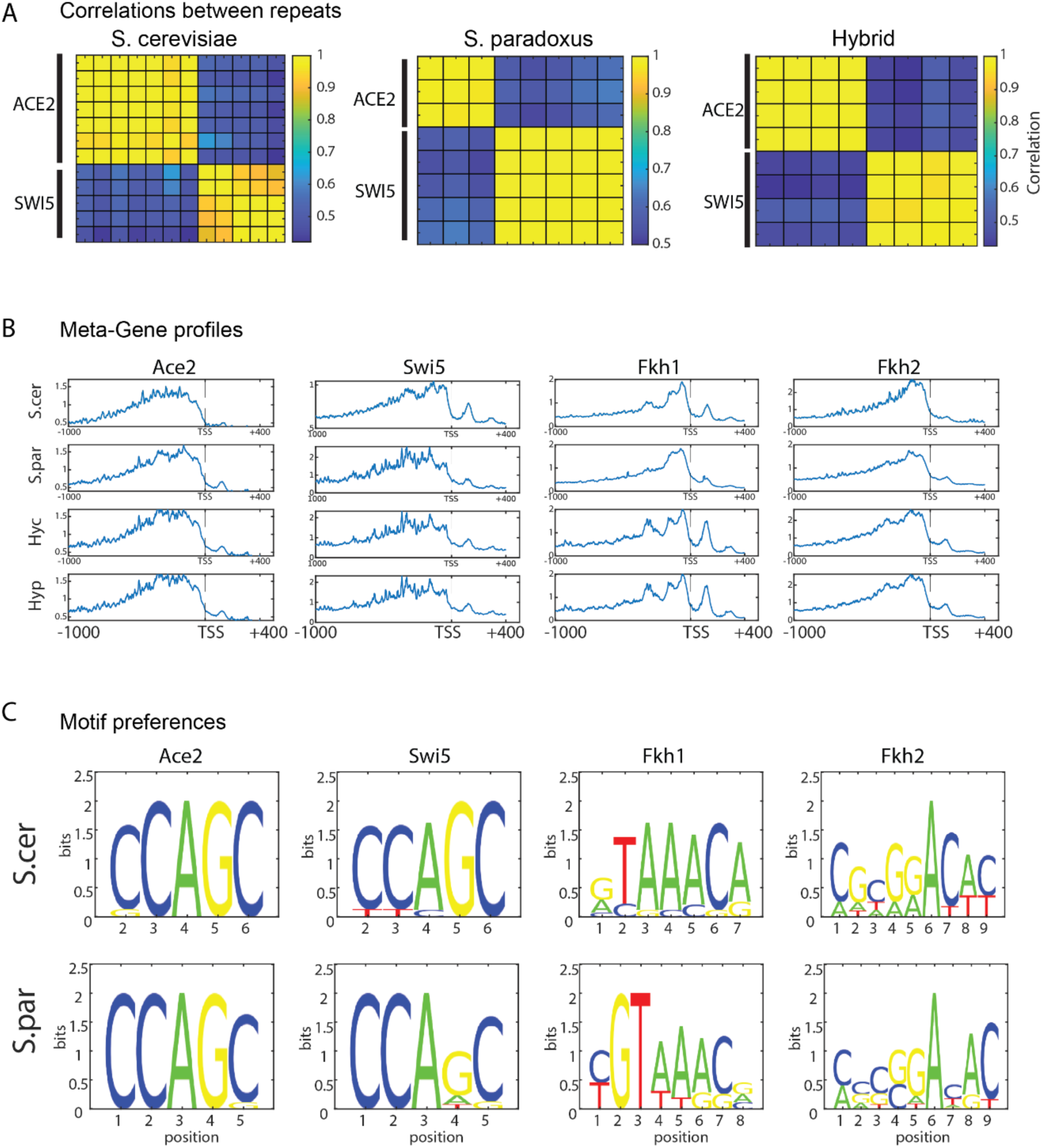
Binding patterns of Ace2, Swi5, Fkh1 and Fkh2. **(A)** Pearson correlation matrices of normalized sum signal on promoters between repeats of Ace2 and Swi5, in S. cerevisiae, S. paradoxus and hybrid (*S. cerevisiae* genome). **(B)** Meta gene profiles of all TFs. All genes were aligned according to their transcription start site (TSS), and the signal was averaged across all genes. (**C)** Sequence logos representing the DNA-motif bound by each factor (See supplementary methods). Ace2, Swi5 and Fkh1 shows binding to their known DNA-motif. Fkh2 doesn’t show a clear logo, though it did get among the top motifs it’s known DNA motif (GTAAACA, not shown). Additional high scoring motifs in Fkh2 might result from interactions with other proteins.

**S5 Fig.**
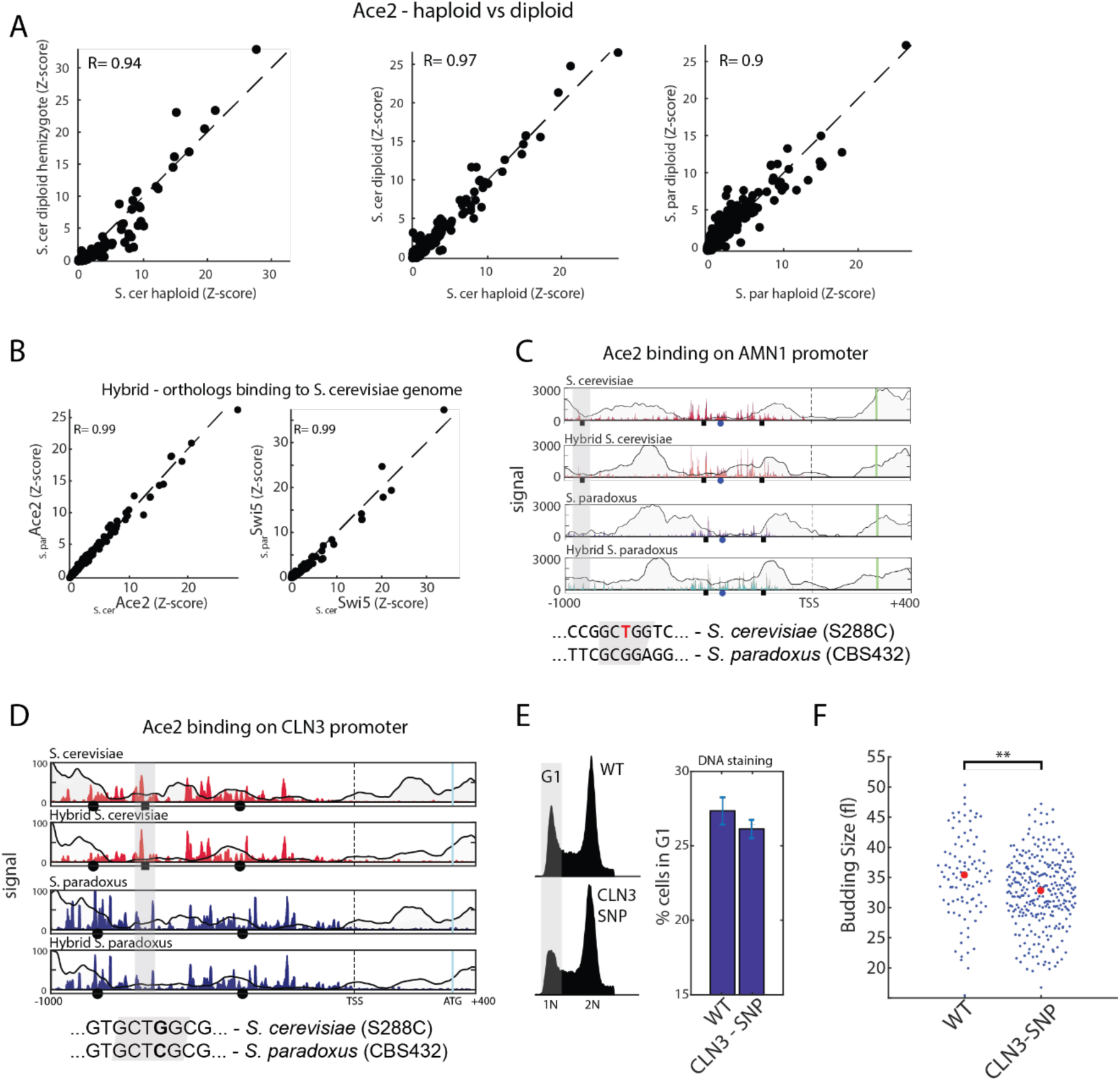
cis and trans effects on Ace2 binding. **(A)** plotted is Ace2 normalized sum of signal on each promoter. Pearson correlation values of top 100 promoters are shown. Left: S. cerevisiae haploid vs S. cerevisiae hemizygote diploid expressing only one copy of ACE2. Middle: S. cerevisiae haploid vs diploid. Right: S. paradoxus haploid vs diploid. **(B)** Normalized sum of signal on each promoter of Ace2 and Swi5 in the hybrid, comparing binding of the two orthologs genes of each factor. **(C)** Ace2 binding on AMN1 promoter. **(D)** Ace2 binding on CLN3 promoter. Shown are the sequence changes leading to loss or gain of Ace2 binding site. **(E)** Left: DNA staining profile of S. cerevisiae strain carrying the mutation of S. paradoxus in CLN3 promoter. Right: quantification of % cells with 1N DNA content. Shown are the mean and standard error of 4 repeats. **(F)** quantification of G1 duration in daughter cells in live microscopy (see methods). Reduced budding size indicates reduces cell size control. Asterisks represent significant p-value (two-sample t-test).

**S6 Fig.**
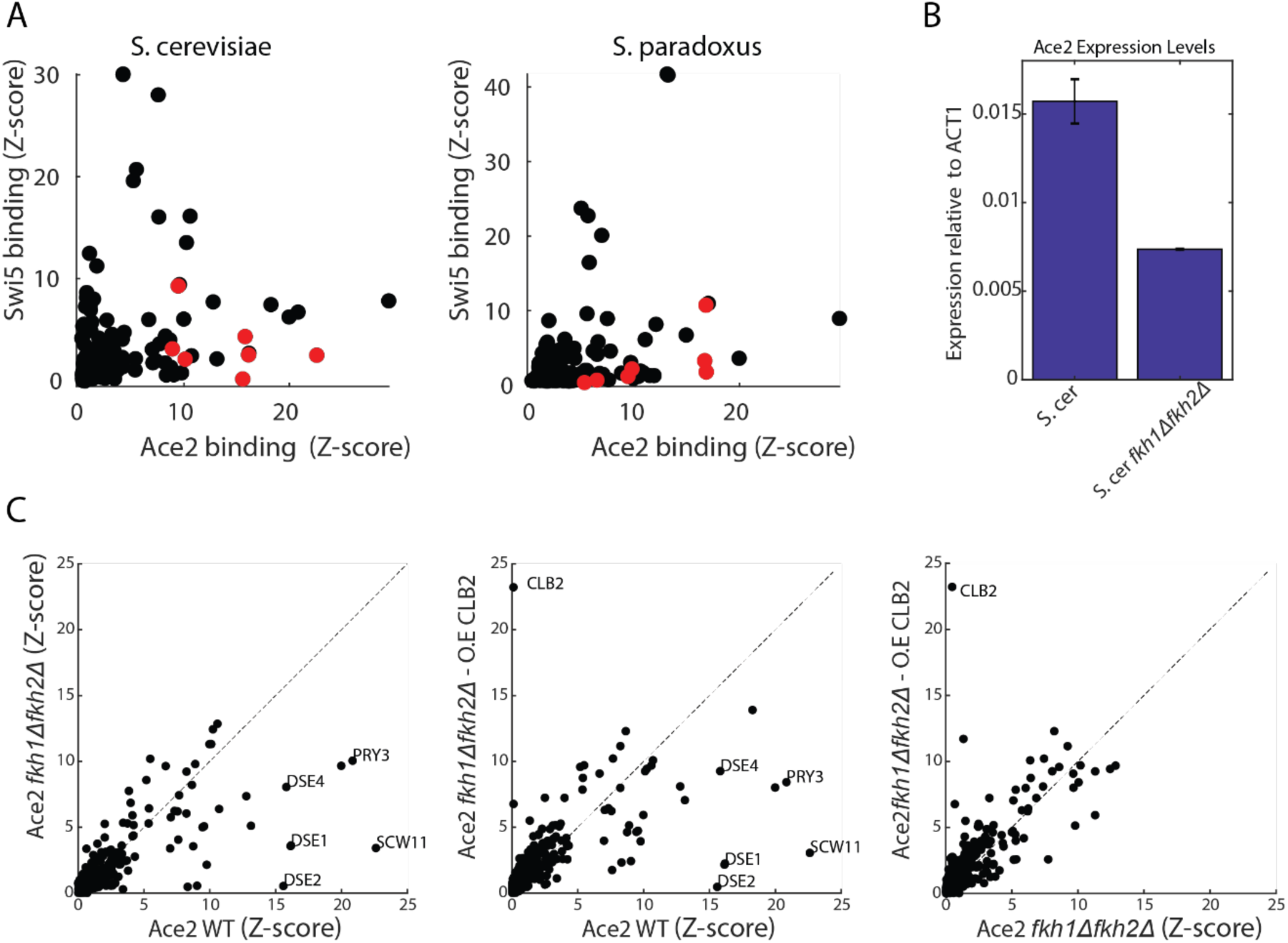
Fkh1 and Fkh2 directly mediates Ace2 binding to cell separation genes. **(A)** sum signal on promoter of Ace2 and Swi5 in S. cerevisiae and S. paradoxus. Red dots indicate cell separation genes. **(B)** ACE2 relative expression levels in WT and in *fkh1*Δ*fkh2*Δ, as measured by RT-qPCR. **(C)** over expression of CLB2 doesn’t restore Ace2 binding to cell separation genes. Shown are sum signal on promoters of Ace2 in WT vs *fkh1*Δ*fkh2*Δ, WT vs *fkh1*Δ*fkh2*Δ CLB2-O.E, and *fkh1*Δ*fkh2*Δ vs *fkh1*Δ*fkh2*Δ CLB2-O.E.

